# Price structure of traded honey, pollen and wax linked to winter survival of a honey bee colony

**DOI:** 10.1101/2024.02.20.581238

**Authors:** Mathieu Kohli

**Affiliations:** Independent

**Author notes:** Contributing authors.

## Abstract

We consider the contribution of both honey and pollen to winter survival of a bee colony in order to assess how valuable both these commodities are from the bees’ perspective. In turn, this leads us to a method to help beekeepers evaluate the marketprice of honey and pollen.

## Introduction

The interaction between *Apis mellifera* honeybees and beekeepers raises the issue of how they can mutually beneficially cooperate? While bees struggle to be resilient through tough weather conditions, beekeepers need to sell honey, wax and pollen at a reasonable price. Here we investigate from a mathematical perspective how a resilient bee colony would price its own honey and other commodities depending on its internal state and on environmental factors.

We start by assuming the environment is a deterministic function of time. Then we allow randomness on the environment to understand how that changes the bees’ perception of the price of honey.

If a beekeeper has this price information in mind, he only needs to add the price of his own labor to compute the price at which he can set his honey. Moreover, since this model focuses on an ideal resilient bee colony, it can be compared to real colonies to discriminate which ones have a high resilience behavior *in the climate in which they live*. Another potential application of the model is the prediction of swarming behavior.

## 1 Value of honey resources

The starting point of our study is the knowledge that a specific flower field or bush or tree in the vicinity of a hive has a limited potential to feed the hive. Inasmuch as the flowers only provide nectar to bees (as opposed to pollen), this potential can be defined as the *net* energy intake gained by the colony *per time unit* as it sends *one* forager bee back and forth between the hive and that resource. We call it the potential efficiency of a forager bee on this nectar resource. It depends on parameters that are identified in [Kohli 2023, Kohli and Anstett]:

- the greater the distance *d* separating the hive and the floral resource, the more energy is spent to transport nectar [Ausseuil et al. 2018] and the longer it takes to travel between these two points. To this, we must add the effect of wind, which is equivalent to a distortion of distances on the geographic map Kohli and Anstett]. Distances can be compared to the maximal distance from the hive *d*_*max*_ at which bees can be found foraging [Beekman and Ratnieks 2000, Greenleaf et al. 2007, Ratnieks and Shackelton 2015].
- The flowers can be highly or scarcely rewarding for bees. We measure this quality through a coefficient *q*_*f*_ that is defined as the energy contained in the stomach of a forager that is full with nectar from flower *f*, to which we substract the energy needed to transform this quantity of nectar into honey through evaporation of superfluous water [Mitchell 2019]. Quantum *q*_*f*_ can be compared to *q*_0_ that corresponds to a reference flower which is the most rewarding flower possible. The flower species also influences the average number of flowers *m*_*f*_ a forager must visit to fill its stomach with nectar, as well as the average time *β*_*f*_ a bee spends on a flower to forage.
- The higher the density *ρ*_*f*_ of flowers, the less energy and time is lost collecting nectar within a small range. Density *ρ*_*f*_ should be compared to a critical density *ρ*_*f,crit*_ below which flowers are too scattered to form a resource.
- Some parameters do not depend on the floral ressource but must be deduced for experiments. Namely *V*, the speed at which foragers travel between the hive and the flowers [Nachtigall et al. 1995], as well as *v*_*hop*_, the average speed at which bees hop from one flower to another. The last relevant parameter that is included in the model we consider is the average time *t*_*hive*_ a forager spends inside the hive between two trips to a floral resource [Fewell et al. 1991, Colin et al. 2022].

Once the parameters that affect the potential efficiency *η*_*f*_ of a forager bee on a nectar resource have been stated, we can express it through the following formula [Kohli 2023, Kohli and Anstett]:

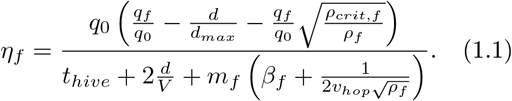

If we assume the bee colony optimizes the use of its foragers on nectar resources, it should send as much foragers as possible to the resource where the foragers have the maximal forager efficiency. If some foragers remain unemployed, they should be affected to the second best resource in terms of efficiency of the bees and so on until no forager is left idle.

To implement the previous process, the knowledge of how many foragers *ϕ*_*f*_ a floral resource can constantly feed is required. It can be derived from the same parameters that were used to express the efficiency of foragers [Kohli 2023, Kohli and Anstett]:

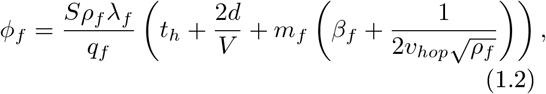

where *S* is the surface of the floral resource and *λ*_*f*_ represents the power offered to foragers by one individual flower [Baude et al. 2016].

## 2 Relative price of pollen compared to honey

To include pollen as a resource in the allocation process previously presented, there is one main problem. Unlike honey, pollen does not provide the energy the hive needs. It provides the matter (amino acids, essential fatty acids, vitamins,…) necessary to form bees [Di Pasquale et al. 2013]. Therefore, the *net gain* for the bee colony brought back by a forager bee is a quantity of matter minus an energy, which can not be directly reduced to one unity. Even if we are able to compare how efficiently bees can forage on two distinct pollen resources, another difficulty follows: how can we compare a nectar resource to a pollen resource? To simultaneously overcome both these challenges, we admit that each day during the foraging season, bees are able to assign a value to both nectar and pollen based on their current needs. The ratio between the value of high quality pollen and the value of honey is the *relative price* of pollen in honey.

To implement this in practice, we replace *q*_*f*_ in formula 1.1 by

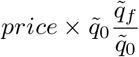

where

- 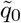 is the maximal mass of pollen that a forager can transport on its legs,
- *price* is the energetic equivalent of the maximal mass of pollen 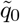 *provided this pollen s of the highest qual ty possible poll n c flower*, in the sense that bees that eat this pollen live the longest (see [Di Pasquale et al. 2013])
- 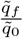 is the ratio between the average lifespan of a bee fed with pollen of flower *f* and the average lifespan of a bee fed with the reference flower [Di Pasquale et al. 2013].

To summarize, a relative price of pollen in honey corresponds to the distribution of foragers on the local floral resources, which in turn corresponds to a certain inflow of nectar and pollen in the hive.

Now the nectar and pollen inflow can be linked to the variation of the reserves of both these commodities by substracting the losses to the inflows. We write the balance sheet for energy as it is stated in [Kohli 2023, Kohli and Anstett]:

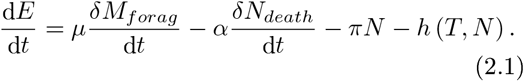

Each term in the previous equation has a specific meaning:

- 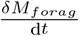 stands for the *net* quantity of honey produced with the nectar inflow entering the hive through foraging. This term can be computed as

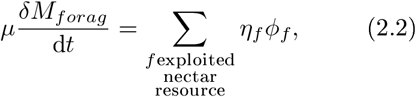

where *ϕ*_*f*_ is computed through 1.2 except for the resource where the foragers are the least efficient, where *ϕ*_*f*_ simply is the number of remaining foragers that are not working on a better resource.
- 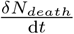 is the death rate in the bee colony. See [Di Pasquale et al. 2013] for a study of this death rate related to the pollen the colony feeds on.
- *πN* represents the “basis metabolism” inside the hive, where *π* stands for the average metabolic rate of a bee that is neither flying nor taking part in the thermoregulation of the hive. [Kovac et al. 2007]
- *h* represents the energy used to thermoregulate the hive Stabentheiner 2021]. It is a function of the ouside temperature and the number of worker bees, but also of the number of males, the number of nurses and the size of the brood.

A similar equation holds for pollen and the number of bees this pollen is transformed into. It has terms equivalent to those in Equation 2.1 except for the term *h* in Equation 2.1 that has no pollinic equivalent.

## 3. Winter survival rate and quantitative state of the colony

At the begining of winter, there are two major factors that depend on the work of the bees during the previous month and that account for the bees’ ability to successfully reach the next spring alive. Those factors are the quantity of honey enclosed in the hive and the number of bees surounding the queen.

Either through a thermodynamic simulation of energy consumption as a function of probable winter conditions outside the hive [Omholt and Lonvik 1986, [Myerscough 1993, Ocko and Mahadevan 2014, Kohli 2023, Kohli and Anstett] or by collecting experimental data [Döcke et al. 2019], one can establish the probabilty 𝕡 (*t* = *fall ⤏winter, N, H*) of surviving through winter if at the time of transition between fall and winter, there are *N* bees and *H* kilograms of honey inside the hive.

At times during the foraging season, in order to describe the quantitative state of the bee colony, it can be relevant to decompose *N* into several components that list the number of bees of different age groups and *H* into an actual honey component and a wax component (since bees process honey to obtain wax) as well as a component of heat accumulated in the form of hive temperature.

We now endeavour to compute, at any time *t* between the begining of spring and the end of fall, 𝕡 (*t, N, H, Envir*). The previous quantity 𝕡 (*t, N, H, Envir*) is defined as the probability to survive the following winter if on day *t*, there are *H* kilograms of honey and *N* bees in the hive, if all the environmental parameters from spring to fall which include flower ressources and weather are known in advance.

The function

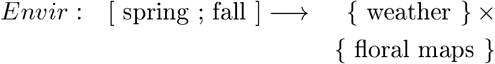

contains all environmental information during the foraging season *including future times* after *t*.

The complete knowledge of the environment between time *t* and the begining of winter implies determinism in the optimal choices of the colony between time *t* and the begining of winter, so that all the randomness comes from winter. And at any time before winter, we can write

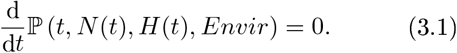

This translates into

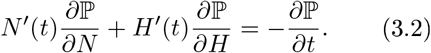

In the previous equation, term 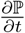 corresponds to the loss of probability of survival in winter that would arise if the hive was somehow “*put onto pause mode*” between time *t* and time *t* + d*t*. The optimization of the survival rate in winter requires to maximize 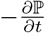 through an optimal choice of foraged resource, which is equivalent to an optimal choice of price:

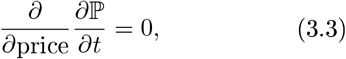

so by Equation 3.2,

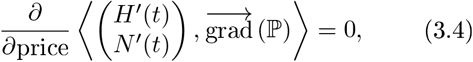

and

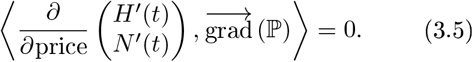

Although 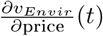 depends on the foraging process *and also on* the internal expenditures of the hive, 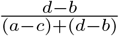 only depends on foraging choices.

The problem of computing 𝕡 for any time *t* during the foraging season can now be solved. Since we assumed that P is known at the begining of winter, we only need to deduce 𝕡 (*t −* d*t*) from 𝕡 (*t*). We start by finding out which price satisfies 3.5, which allows us to discover *H*^*′*^(*t*) and *N*^*′*^(*t*). Then we use 3.2 that leads us to 𝕡 (*t −* d*t*).

## 4 Randomness in the environment

Of course, the future is not fixed and there are different scenarios *Envir* that can play out with different probabilities. We denote by 𝕊 the probability on the set of environmental conditions.

To understand what can happen when there are different paths leading from time *t* to the begining of winter, let us assume that there are just two different functions *Envir*1 and *Envir*2 that describe the same local conditions up to time *t*_*r*_ but diverge at time *t*_*r*_. The hive must *bet* on *Envir*1 or *Envir*2 and follow the optimal path in the state space (*H, N*) up to time *t*_*r*_ + d*t* when it will be able to reassess if it did the correct choice or not. But if there is not only one but several hives in the same state (*H, N*), some will choose *Envir*1 and others will choose *Envir*2, which means that the state (*H, N*) will split into two states that increasingly diverge until time *t*_*r*_ +d*t*. To know in which proportions both choices are made, we refer to Appendix A. Now at time *t*_*r*_ + d*t*, all hives realize which choice was right, for example *Envir*1, and both splited states follow the optimal path in the state space (*H, N*) knowing they are in *Envir*1 from there on. This leads to a splited state at the begining of winter.

By knowing the proportion of hives in each state and the probability to survive winter in each case, we deduce the global probability to survive winter along the environmental conditions *Envir*1, *ramified at time t*_*r*_ if at time *t* we are in the state (*H, N*) that we denote by

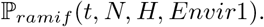

The previous notation stands for any ramification structure, even if it is more complex than Just one ramification at time *t*_*r*_. To replace a simple state in the (*H, N*) *−* space by any splited state, we introduce probability densities on the (*H, N*) *−* space. A simple state then becomes a Dirac density.

Now at time *t*, the actual probability to survive winter can be written as

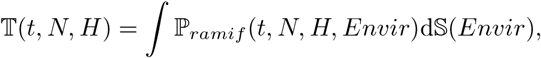

where it is understood that d𝕊 is conditioned to the knowledge of what happend in the season before time *t*.

## 5. Absolute prices

Now that the meaning of a relative price of pollen in honey has been clarified, we are ready to define an absolute price of both honey and pollen in a currency such as dollars, euros,… Indeed, let us assume a beekeeper has only one hive and the market price of a bee swarm is for example 200 euros. This is the price a beekeeper would have to pay to maintain his activity if he is *certain* that his bee colony will not survive winter. Conversely, if the beekeeper is *certain* that his hive will live through winter, that saves him 200 euros. This means that the value of 𝕋 = 1 is 200 euros.

This means that the price of a kilogram of honey is given by

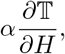

where we took for example *α* = 200 euros.

This price in turn translates into an equivalence between currency and pollen through the relative price of pollen in honey.

Let us notice that wax prices can also be computed. The production of wax does not compete with the production of honey or pollen. Indeed, it is when they are between 5 and 20 days old that bees produce wax and build wax structures, while they usually start foraging after they are three weeks old Clement and Le Conte et al. 2018]. The production of 1g of wax requires around 8.4g of honey according to [Rodney and Purdy 2020] and since wax production does not interfere with any other production, the price of wax is simply 8.4 as much as the price of honey.

The best interest of both the beekeeper and the bees is that the honey, wax or pollen harvests happen at the moment when these commodities are at their lowest price. The beekeeper will add the cost of his own labour to this price when he sells his honey.

## 6 Swarming conditions

For a hive to form a swarm, the probability that both halfs of the initial colony die before the begining of the next spring should be inferior to the probability that the initial colony dies before that same moment. This translates into

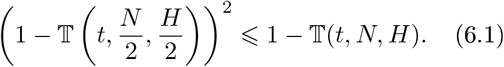

Here, we did not take into account the dissymmetry between the old queen that leaves with the danger of wandering and the new queen that stays inside the hive with all the wax structure already built.

Let us notice that the beekeeper has one major advantage over the bee colony when it comes to computing 𝕋: he has access to meteorological predictions for the next few days. At a pivotal moment of the year such as swarming period, that means that he can anticipate swarming behavior.

## 7 Hornets

Bees are to hornets what pollen is to bees: food to feed their larva. From that perspective, we can apply the pricing results that apply to honey bees to hornets similarily. This can offer insight to find out if hornet traps near beehives makes hunting bees an unsustainable activity or not. The value of lost hornets must be considered *pro rata temporis* of their remaining life expectancy and should be compared to the number of hornet larvae that can be fed with the bees that get captured around the hive.

## Appendix A

**Betting on two opposite teams simultaneously**

Let us assume a player is given the choice to bet between two outcomes.

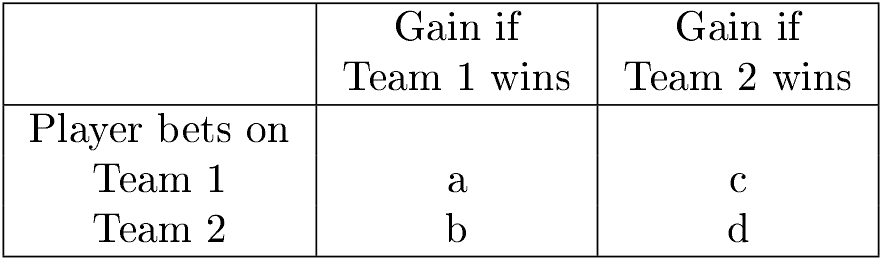

|  | Gain if<br>Team 1 wins | Gain if<br>Team 2 wins |
| --- | --- | --- |
| Player bets on<br>Team 1 | a | c |
| Team 2 | b | d |

To be consistent, we impose *b < a, c < d*.

If instead of betting all on one team, the player decides to bet 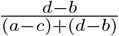 on Team 1 and 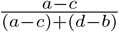 on Team 2 then the player knows he will always win 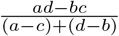. This strategy is the most resilient one, as it brings at its maximum the amount that is won with certainty.

If it so happens that the amount that should be bet on one team is negative, which is the case if and only if *a < c* or *d < b*, this leads to betting entirely on the opposite team. This corresponds to the “preparing for the worst, hoping for the best” strategy.

In the case we are studying, we translate this using the dictionary

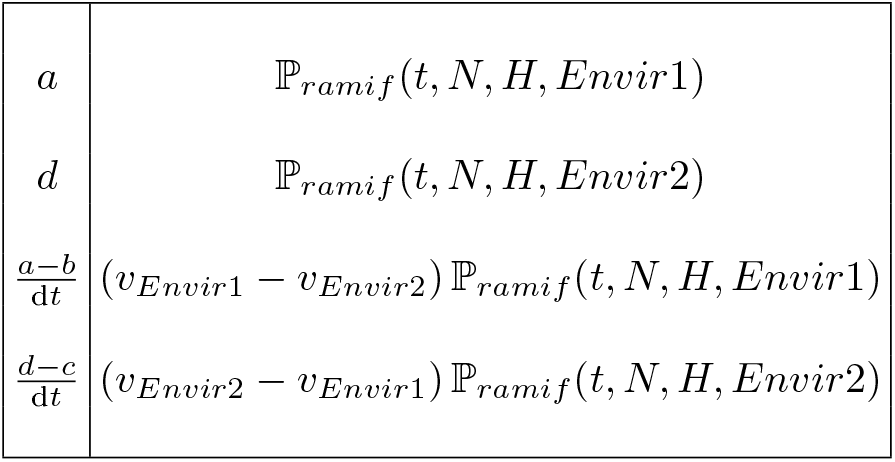

Here, d*t* denotes the time it takes for the hive to realize which of the two environmental scenarios actually is taking place. In the case where *c < a* and *b < d*, if d*t* tends to zero, the gain that can be assured at the ramification point is the translation of 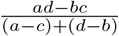 :

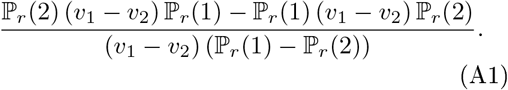

On the other hand, if *a < c* or *d < b*, randomness must still be taken into account and the hive bets on the worst case scenario.

